# Neural Dynamics Underlying False Alarms in Extrastriate Cortex

**DOI:** 10.1101/2024.09.06.611738

**Authors:** Bikash Sahoo, Adam C. Snyder

**Affiliations:** Brain & Cognitive Sciences, University of Rochester, Rochester, NY 14627, USA

## Abstract

The unfolding of neural population activity can be approximated as a dynamical system. Stability in the latent dynamics that characterize neural population activity has been linked with consistency in animal behavior, such as motor control or value-based decision-making. However, whether similar dynamics characterize perceptual activity and decision-making in the visual cortex is not well understood. To test this, we recorded V4 populations in monkeys engaged in a non-match-to-sample visual change-detection task that required sustained engagement. We measured how the stability in the latent dynamics in V4 might affect monkeys’ perceptual behavior. Specifically, we reasoned that unstable sensory neural activity around dynamic attractor boundaries may make animals susceptible to taking incorrect actions when withholding action would have been correct (“false alarms”). We made three key discoveries: 1) greater stability was associated with longer trial sequences; 2) false alarm rate decreased (and reaction times slowed) when neural dynamics were more stable; and, 3) low stability predicted false alarms on a single-trial level, and this relationship depended on the elapsed time during the trial, consistent with the latent neural state approaching an attractor boundary. Our results suggest the same outward false alarm behavior can be attributed to two different potential strategies that can be disambiguated by examining neural stability: 1) premeditated false alarms that might lead to greater stability in population dynamics and faster reaction time and 2) false alarms due to unstable sensory activity consistent with misperception.

## Introduction

In a dynamical system, the future system state can be predicted from the current system state. Considering a neural population (many neurons within a brain area), this means that if we know the activity levels of a sample of neurons at one time (the “initial condition”), we can roughly predict the activity levels of those neurons some time into the future. This doesn’t necessarily mean an unchanging pattern of activity will persist into the future; rather, the pattern of activity may evolve according to a set of rules (the rules are not usually explicitly known, but we can infer such rules exist from the predictability of the dynamics). The longer into the future an accurate prediction about system states can be made, the greater the dynamic stability. In some cases, many different initial conditions may converge onto a common state in the future. That common state is said to be an “attractor” for the system, and the set of initial conditions leading to that attractor form the “basin of attraction”. An attractor can be a fixed, unchanging state, or it can be a sequence of states, termed an “attractor cycle”.

Attractor cycles provide an advantageous mechanism for guiding behavior because they are robust to perturbation. That is, small deviations of the system state from the attractor cycle due to noise or outside influence will remain within the basin of attraction and be drawn back to the attractor cycle at some point in the future (in the near-future for a dynamically stable system, in the further-future for a less-stable one). For example, Li *et al*. (2016) tasked mice to locate objects by whisking, then during movement preparation the experimenters optogenetically perturbed large portions of the premotor cortical network. They found that the premotor network as a whole rapidly compensated for this perturbation consistent with a dynamic attractor, leading to motor behavior being largely unaffected. The dynamical systems framework for behavior has benefited the study of motor control (Shenoy *et al*., 2013), but the extent to which such dynamic principles guide perceptual processing is much less understood.

For example, when on a long road trip on a highway, much of the task is to remain vigilant, and, having determined that no hazards have arisen, take no action. This perceptual decision-making loop leading to inaction is then repeated for hundreds of kilometers (hopefully). Many tasks share this pattern of vigilant perception and inaction. However, it is common to spontaneously break out of this deliberative loop, leading to a suboptimal break in concentration or impulsive action (“false alarm”). We hypothesized that such perceptual decision-making loops are guided by dynamical systems, much as for motor control, and that false alarms may be caused, at least in part, by a weakening of dynamic stability that dictates the evolution of system states. Specifically, we theorized that the neural state in macaque extrastriate visual area V4 traverses a dynamic attractor on each iteration of the perception-decision loop, and that periods preceding false alarms would be characterized by slower attraction towards that attractor, consistent with weakened dynamic stability.

## Results

We recorded local field potentials (LFPs) from area V4 of two adult male macaques engaged in a non-match-to-sample grating orientation change-detection task (Figure 1a). Separate analyses of other data from these experiments have been previously reported (Snyder *et al*., 2018; Cowley *et al*., 2020; Snyder *et al*., 2021; Umakantha *et al*., 2021; Johnston *et al*., 2022; Sachse & Snyder, 2023). The task included a spatial attention manipulation across blocks of trials, and our previous reports concerned spatial attention effects and did not specifically analyze false alarm behaviors. Because animals overwhelmingly directed false alarms towards the spatially attended stimulus (89.8% for M1, 90.1% for M2), we did not analyze false alarm behavior in the context of spatial attention for this report. In general, monkeys performed the task well, correctly discriminating targets with an orientation change from repeated standard stimuli (i.e., no change; *d*^*′*^ = 1.24 (0.13) [*mean* (*SD*)] for M1, *d*^*′*^ = 0.67 (0.29) for M2). However, performance was not perfect, and monkeys incorrectly made saccades to standard stimuli (*false alarms*) at a moderate rate; 0.25 (0.06) false alarms per standard stimulus presentation for M1, 0.37 (0.11) false alarms per standard stimulus presentation for M2. One feature of our task was that trials could contain several repeated standard stimuli before the presentation of a target, requiring the animals to perform a cognitive “loop” of deliberative inaction (number of repetitions *N* ≤ 19, mostly 1–3 repetitions per trial, supp. fig. 1). We reasoned that this repetitive task structure might be particularly well-suited for revealing the nature of the relationship between stable behavior and neural population dynamics. To build a cohesive and mechanistic explanation of the false alarm behavior, we studied three types of relationships, 1) the correlation between animals’ false alarm rate and time-lapsed since the trial onset, i.e. non-target (standard) stimuli repetition number, 2) the correlation between estimates of stability of neural trajectories (i.e. stability index) and stimuli repetition number, and 3) the correlation between stability index and reaction time.

**Figure 1.**
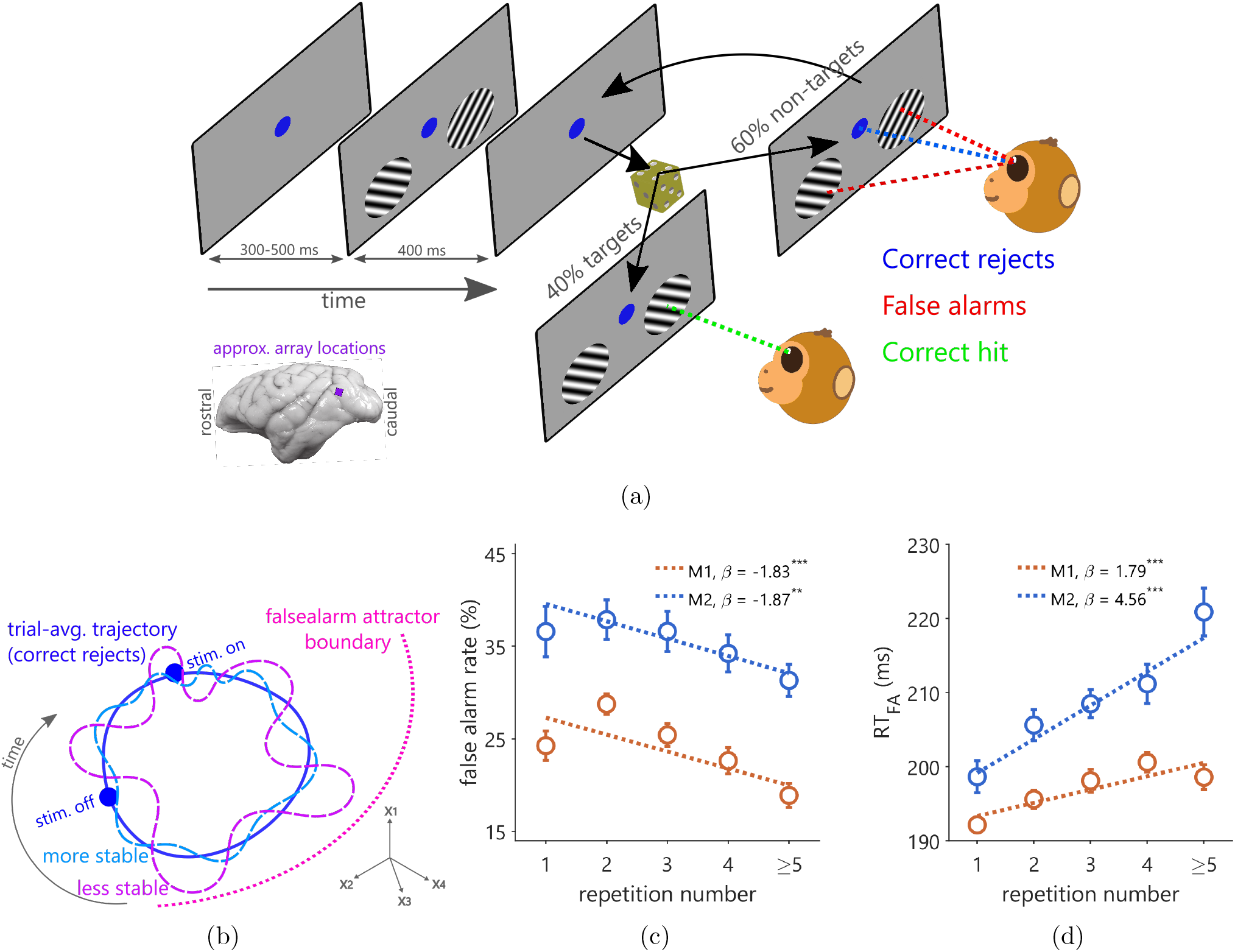
Experimental paradigm and behavioral results. (a) Non-match-to-sample change detection task; each trial sequence contained several repetitions of the sample stimuli (1st set of stimuli or non-targets) before an orientation-changed stimulus, i.e. target, appeared in either location. The monkey was required to saccade to the target stimulus. (b) Schematic of our hypothesis: state-space illustration of sequences of neural activity states (neural trajectories) for a trial sequence with 2 repetitions of the non-target stimuli where the monkey correctly withheld saccade. One neural trajectory is more stable (cyan) and therefore less likely to escape the attractor boundary (magenta) and lead to a false alarm compared to the other, less stable neural trajectory (purple). (c) False alarm rate decreased as stimuli repetitions increased (inset shows regression coefficients and corresponding statistical significance of curve fits between %*FA* and repetition number for individual monkeys; ** : *p* ≤ 0.01, *** : *p* ≤ 0.001). (d) FA reaction time slowed as stimuli repetitions increased. That is, when monkeys experienced longer sequences of non-target stimuli, they were less likely to commit false alarms and were also slower to act if/when they did so.

### False alarm behavior improved with repetition number

We examined how the animals’ false alarm behavior was related to the number of such deliberative loops that had been performed on each trial (i.e., the “repetition number”). Since trials with more such loops would lengthen the time the animal needed to engage with the task before being rewarded, one potentially relevant phenomenon affecting monkeys’ behavior could have been so-called “delay discounting”: across various animal species, such as primates, rats, birds, etc., it has been commonly observed that the subjective value of a future reward reduces with the delay before it is availed (Vanderveldt *et al*., 2016; Hwang *et al*., 2009). For similar magnitudes of rewards, animals prefer actions that lead to immediate reward compared to delayed ones. This may lead the monkey to make more false alarms when the number of stimuli repetitions becomes higher thereby deteriorating behavioral outcomes in the task. However, in our experiment, to alleviate the effect of such a discounting phenomenon and to equate motivation across variable trial durations, we increased the reward amount exponentially as the trial duration increased.

Beyond the delay-discounting mechanism that we took efforts to mitigate, we considered two potential cognitive mechanisms whereby false alarm behavior and repetition number could be associated. One mechanism could be that repeating a behavior causes it to be more likely to be executed again in the future, through, e.g., Hebbian plasticity mechanisms (Hebb, 1949). Another mechanism could be that if the animal is in a state indisposed to false alarms, this would lead to the animal “lasting” longer into the trial and therefore experiencing more stimulus repetitions (i.e., a survivorship effect). Both of these hypothetical mechanisms predict an inverse association between false alarm rate and repetition number, and a direct association between false alarm reaction time (*RT*_*F A*_) and repetition number. To test for such associations, we calculated the monkeys’ false alarm rate and *RT*_*F A*_ as a function of repetition number in each session. We considered a decrease in false alarm rate (%*FA*) and a slowing in *RT*_*F A*_ as a signature of behavioral improvement. Our results concurred with these predictions (fig. 1c and 1d; Spearman’s rank correlation between false alarm rate and repetition number *ρ* = −0.34 (*p* = 0.0001) for M1 across sessions (*N* = 24) and *ρ* = −0.23 (*p* = 0.0143) for M2 (*N* = 23); rank correlation between false alarm RT and repetition number *ρ* = 0.35 (*p* = 0.0001) for M1 and *ρ* = 0.51 (*p* = 7.520 × 10^*−*9^) for M2. Estimates from a generalized linear mixed-effect model with session number as the random-effect term (equations 3 and 4 in Methods) produced similar results; for false alarm rate, *β*_*repetition*_ = −0.0090 (*P <* 0.001, 95% *CI* = [−0.0117, − 0.0060], *N* = 24) for M1 and *β*_*repetition*_ = −0.0078 (*P <* 0.001, 95% *CI* = [−0.0116, − 0.0034], *N* = 23) for M2. Similarly for false alarm RT (equations 5a and 5b), *β*_*repetition*_ = 0.0017 (*P <* 0.001, 95% *CI* = [0.0012, 0.0021], *N* = 24) for M1 and *β*_*repetition*_ = 0.0044 (*P <* 0.001, 95% *CI* = [0.0040, 0.0051], *N* = 23) for M2. This pattern of improved (slower and less common) false alarm behavior as repetition number increased suggests that our delay-adjusted reward schedule successfully countermanded subjective reward discounts, and suggests the potential for our additional hypothesized mechanisms linking dynamic neural stability to false alarm behavior, such as reinforcement through plasticity or survivorship effects.

### Variance in LFPs explained by repetition number

Next, we sought to understand the relationship between stimulus repetition number and neural activity. First, we estimated how much of the variance in the LFPs can be explained by repetition number. We performed demixed principal components analysis (dPCA) on the LFP data and considered attention condition, stimulus orientation presented in the receptive field, and repetition number as three marginalization factors in the analysis (Kobak *et al*., 2016). The average variance contribution of the principal component for cue was 0.10% (*SEM* 0.02%), for stimulus orientation 2.06% (0.31%) and for repetition number 0.21% (0.05%). The condition independent variance explained in the LFPs was estimated to be 55.24% (3.01%). Both cue and stimulus orientation could be more reliably decoded from LFPs compared to repetition number. Cross-validated discrimination accuracies of cue, stimulus orientation and repetition number during stimulus presentation period were 54.24% (*SEM* 0.45%, *chance* 50%, *p* = 2.046 × 10^*−*149^), 75.39% (*SEM* 2.69%, *chance* 50%, *p* = 1.648 × 10^*−*113^), and 27.16% (*SEM* 0.26%, *chance* 25%, *p* = 8.927 × 10^*−*147^) respectively. Thus, the LFPs reliably encoded key experimental variables.

### Repetition number predicted stability in population dynamics

We estimated the dimensions in the LFP activity subspace that maximized the stimulus signal-to-noise ratio. We used a generalized eigenvalue method to identify the dimensions that maximally explained activity elicited as a response to stimulus while minimizing the residual noise (de Cheveigné & Parra, 2014). The trial-averaged response in this subspace we termed as the “*stable trajectory* “, our estimate of the hypothesized limit cycle attractor (fig. 2b). One aspect of a stable system is that when its stable state/trajectory is perturbed, it subsequently acts against the perturbation and converges onto its prior steady state/trajectory. Our estimation of stability hinges on this logic. At each time point in the trajectory, we estimated the projection of the change in residual activity at the immediate future time point along the direction of the original trajectory, which is the *“pulling force”* towards the trajectory when a deviation occurs away from it (fig. 2b). We computed a metric for estimating stability in neural dynamics i.e. stability index as the net “*pulling force*” towards the stable trajectory over a time period. For each repetition number we estimated the average pulling force at each time point and the net stability during the stimulus presentation period. We found that stability index estimates and repetition number were positively correlated (fig. 3), suggesting the stable trajectory became more attractive with each successive stimulus presentation (Spearman’s *ρ* = 0.0164, *z* = 0.0164, *p* = 0.0373 for M1 (*N* = 24 sessions) and *ρ* = 0.0471, *z* = 0.0472, *p* = 4.577 × 10^*−*6^ for M2 (*N* = 23)).

**Figure 2.**
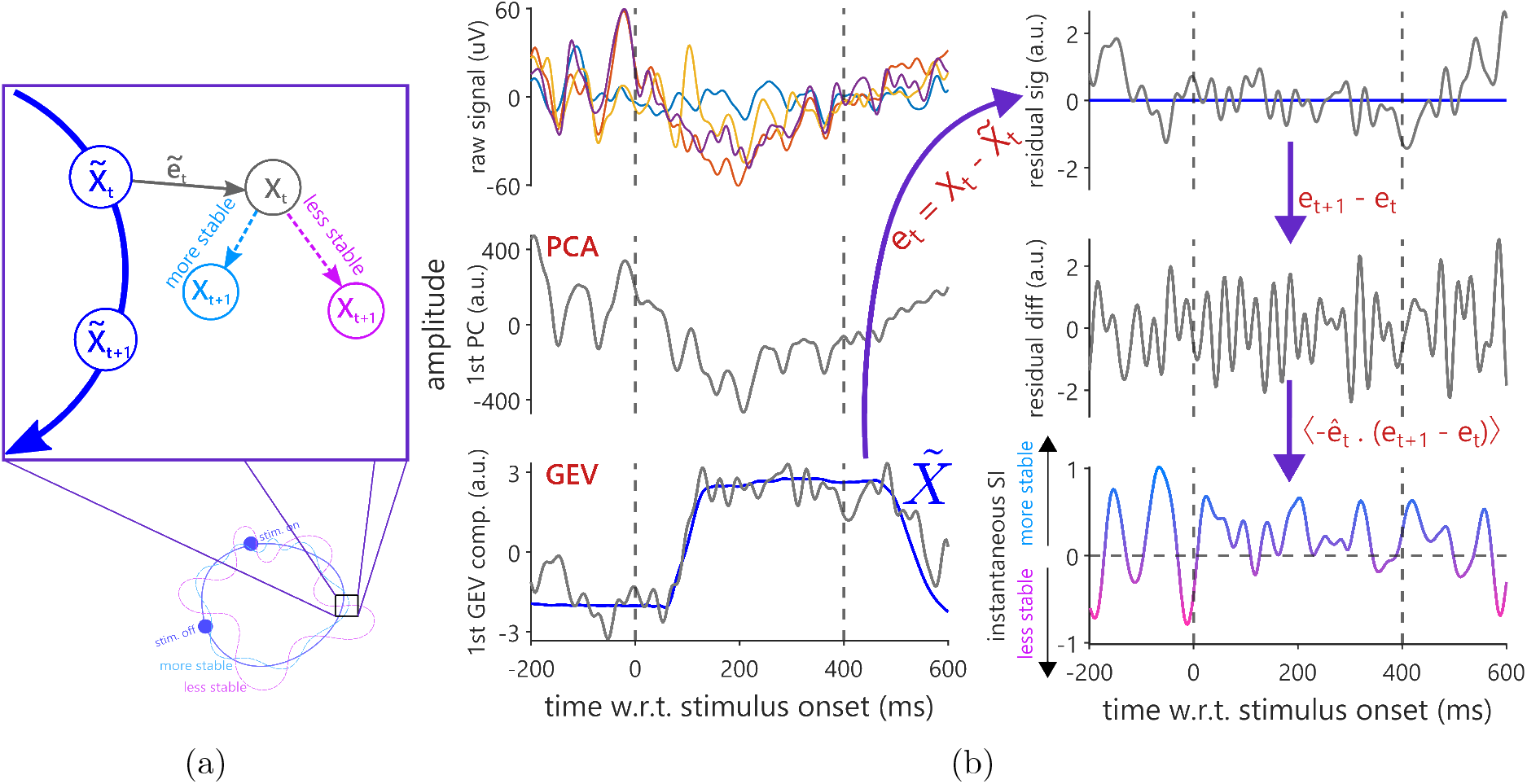
Estimation of stability of neural trajectories and the relationship between stability and stimuli repetitions. (a) Schematic of computing stability at a given time point *t*, as the amount of change in perturbation in the direction of the stable trajectory (blue). (b) Left-top: raw voltage traces of five randomly chosen channels for a single representative trial; left-middle: time series corresponding to the same single trial for the top principal component; left-bottom: time series for the same trial after dimensionality reduction was done using GEV to maximally separate SNR i.e. signal common across all trials versus residual noise, showing GEV can faithfully separate stimulus-evoked response (step-like dynamics) from noise. Right-top: residual activity in the trajectory corresponding to maximum SNR estimated after subtracting out the trial-averaged signal; right-middle: instantaneous changes in residual activity; right-bottom: instantaneous estimates of stability index as a normalized product between residual activity and its instantaneous changes. Purple arrows indicate the analysis pipeline. Inset texts in red show the computations performed.

**Figure 3.**
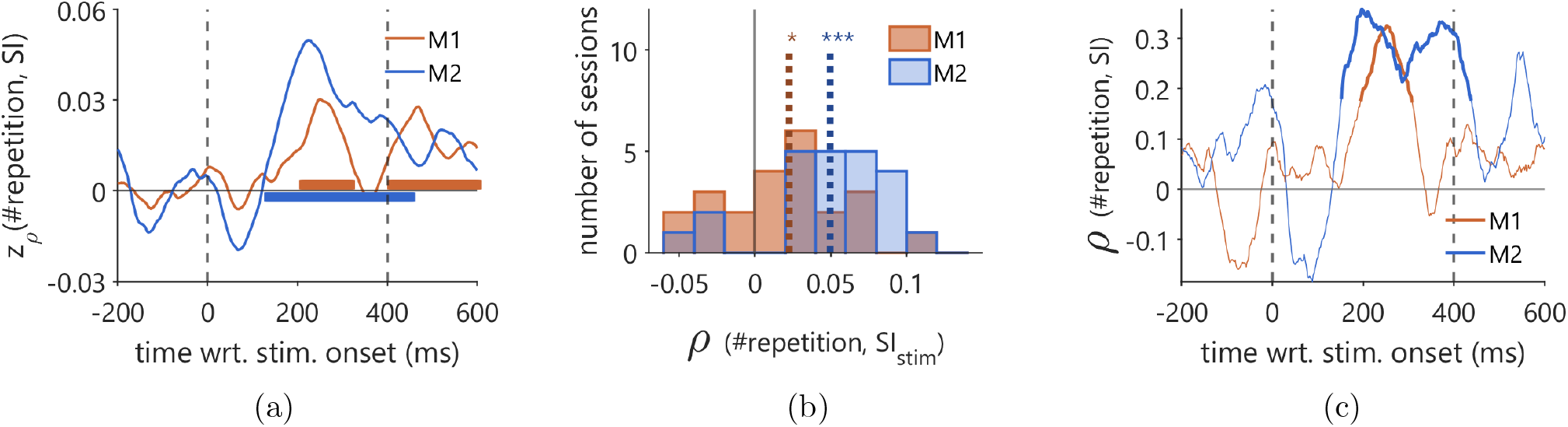
Neural activity during the stimulus response grew more dynamically stable as stimuli repetition increased. (a) Session-averaged time series of Fisher-transformed correlation (*z*_*ρ*_) values between repetition number and stability index, computed in each session. Thick lines indicate periods where correlation significantly differed from zero across sessions (two-tailed cluster-based permutation test, *p* ≤ 0.05). (b) Histogram of individual correlation values computed between SI averaged over a time window 100-400ms after stimuli onset and repetition number in a session. (c) Conventions are the same as (a) but, the correlation is computed between repetition number and *SI* across sessions after averaging *SI* values for each repetition number within a session. Thick lines indicate statistically significant correlation values (two-tailed cluster-based permutation test after converting *ρ* to *t*-score using 20, *p* ≤ 0.05).

### Reaction time slowed, and *FA* likelihood decreased, with stability in the neural dynamics

We reasoned that trials with higher estimates of stability index would experience a greater pulling force towards the trial-averaged trajectory (i.e. corresponding to correctly fixated trials), thereby would be less likely to drift away to other parts of the neural state space. In other words, trials with a higher stability index would be less likely to lead to false alarms. We tested this by fitting generalized linear models (GLMs) between neural stability estimates and behavioral trial outcomes for no-change stimuli presentations (i.e., correctly continued fixation versus false alarm saccades). In line with our prediction, we found stability estimates were negatively related to false alarm occurrence (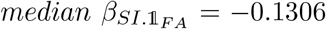, left-tailed *t*_23_ = −2.05, *p* = 0.0258, *N* = 24 for M1, and 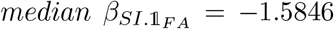, left-tailed *t*_22_ = −12.24, *p* = 1.364 × 10^*−*11^, *N* = 23 for M2). This suggests that the attractive strength of dynamic stability in V4 population activity can “make or break” the animals’ false alarm behavior in the task.

In addition to the previous binary result, one more nuanced prediction of the dynamic stability framework is that, given that a saccade *does* occur, that action should take more time to escape from the deliberative loop of action inhibition when neural activity is more stable, because it has to overcome a stronger attractive force. In this scenario, reaction time will be positively correlated with the stability of the neural system, i.e. the more stable the system is the slower the reaction time will be. Importantly, this reasoning applies not only to the reaction time for false alarms, but also for correct saccades. To test this prediction, we analyzed the monkeys’ LFP aligned to saccade onset in each trial. We estimated the stability index for each trial in a session, and computed the partial correlation between stability index and RT within a session after accounting for the variation due to stimuli repetition number (fig. 4a and 4b). RT was positively correlated with stability index for both false alarms (mean Spearman’s *ρ* = 0.1153, *z* = 0.1167, *t*_23_ = 6.23, *p* = 2.360 × 10^*−*6^ for M1 (*N* = 24 sessions) and *ρ* = 0.2411, *z* = 0.2475, *t*_22_ = 14.72, *p* = 7.189 × 10^*−*13^ for M2 (*N* = 23 sessions), and for correct saccades (fig. 4c and 4d, *ρ* = 0.0840, *z* = 0.0850, *t*_23_ = 4.40, *p* = 0.0002 for M1 and *ρ* = 0.1835, *z* = 0.1876, *t*_22_ = 9.12, *p* = 6.225 × 10^*−*9^ for M2). The consistency in this relationship across different stimulus contexts i.e. being a target or non-target, suggests it to be a generic contributor to saccade behavior.

### Spontaneous dynamic stability was inversely related to strength of visual evoked responses

Under the dynamic attractor framework, dynamic (in)stability is related to the sensitivity of the neural population to perturbation. For a sensory population, this suggests that stimulus input arriving during periods of relative dynamic instability should be associated with more robust evoked responses compared to stimulus input arriving during periods of relative dynamic stability. To test this, we divided trials into quantile bins based on the average SI during the prestimulus spontaneous activity and then quantified the overall magnitude of the evoked response as the global field power (standard deviation of voltage across the electrode array) during the stimulus-response for each bin. Confirming our prediction, we found that prestimulus dynamic stability was significantly negatively correlated with the magnitude of the evoked response to the stimulus (fig. 5). This suggests that dynamic stability may index periods of relatively stronger or weaker visual sensitivity within V4 populations.

### Low and high stability were associated with distinct types of false alarm behavior

Because we found that the V4 population had different visual evoked response magnitudes depending on whether the spontaneous activity was especially stable or unstable, we asked whether dynamic stability might reflect differences in the animals’ engagement with visual processing. For example, a period of high stability might indicate the animal is withdrawn from (or insensitive to) the visual task, while a period of low stability might indicate the animal is engaged in (or sensitive to) it. These different regimes of task-engagement or task-sensitivity would likely be associated with different “types” of false alarm behavior. In the task-engaged/sensitive state, the animal may false alarm because they believed that they perceived a target when in fact no target had been presented (a “misperception”). In the disengaged state, the visual stimulus is unlikely to trigger a false alarm, but rather the animal may decide to make a false alarm for non-sensory reasons, such as impatience with the task (a “premeditated” false alarm). Thus, the difference between these two types of false alarms could be expected to relate to the detailed structure of the task and the timing of visual stimuli, as well as the dynamic stability of V4. In both monkeys, false alarm rate and reaction time covaried with the ISI, i.e., time elapsed since the previous stimulus, during which the monkey was required to maintain fixation. *FA* rate increased, and *RT*_*F A*_ decreased, with an increase in ISI. The trends in these relationships are in contrast with *SI*, where *FA* rate decreased and *RT*_*F A*_ increased, with an increase in SI, which itself increased with ISI (supp. fig. 2). We sought to evaluate if these relationships are interdependent, i.e. if the relationship between ISI and *RT*_*F A*_ varied with an increase or decrease in SI, and whether differences in these relationships could be explained in terms of “premeditated” versus “misperception” false alarms.

### Enhanced stability was associated with “premeditated” false alarms

We divided trials into bins of different combinations of ISI and SI, and estimated *FA* rate and *RT*_*F A*_ in each bin. Specifically, we binned in 12 quantiles of ISI and 8 quantiles of SI, for a total of 12 × 8 = 96 bins of trials covering all combinations. In general, *RT*_*F A*_ was inversely related to ISI across all SI bins (fig. 6). That is, animals were faster to false alarm to a stimulus if they had waited a relatively longer time since the previous stimulus. However, the slope of this relationship became steeper with the increase in SI (fig. 6d; SI below median: *β*_*RT*_ = −0.8895, *t*_2178_ = −7.76, *p* = 1.298 × 10^*−*14^, 95% *CI* = [−1.1143, −0.6647]) and SI above median: *β*_*RT*_ = −1.8070, *t*_2031_ = −12.51, *p* = 1.173×10^*−*34^, 95% *CI* = [−2.0902, −1.5238]). That is, when V4 activity had high dynamic stability, there was an especially strong inverse relationship between ISI and false alarm RT. One simple explanation for such an inverse relationship between ISI and RT is that on these trials the animals were actually trying to “time” their saccades relative to the offset of the previous stimulus in the sequence, rather than reacting to the onset of the current stimulus, because the ISI is the time between the current stimulus (which is not known to the monkey in advance) and the time of the previous stimulus (which is). In other words, the monkeys had made up their mind before the stimulus onset that they were going to make a saccade no matter what (a “premeditated” saccade) and were using the previous stimulus offset in order to anticipate the correct timing. To more directly test this idea, we calculated RTs aligned to the time of the previous stimulus offset. If monkeys were trying to time their responses relative to these previous stimuli, then their RT distributions would have a smaller variance when aligned to the previous stimulus compared to when RTs were aligned to the current one. In contrast, if monkeys were reacting to the current stimulus, then their RT distributions would have a smaller variance when aligned to the current stimulus compared to when aligned to the previous one. We found that for trials with high dynamic stability, RTs were better explained by the previous stimulus than the current one (suggesting premeditated saccades), whereas the converse was true for trials with low dynamic stability (suggesting reactive saccades; fig. 6e).

For a more detailed look at the relative effects of SI and ISI, we divided our space of SI versus ISI into four quadrants (fig. 6a), representing different binary combinations of high and low stability and long and short ISIs (i.e., above or below the median), and compared *RT*_*F A*_ between quadrants. In trials corresponding to both higher ISI and SI, *RT*_*F A*_ (↑ISI↑SI quadrant) was significantly faster compared to trials with lower ISI (fig. 6b; ↓ISI↑SI and ↓ISI↓SI; *t*_139_ = 3.88, *p* = 0.0002). Only for higher ISI, an increase in SI did not translate to a decrease in *RT*_*F A*_ (↑ISI↑SI and ↑ISI↓SI: *t*_46_ = 0.63, *p* = 0.5317).

To summarize the findings thus far: in general, dynamic neural stability was inversely related to the prevalence and speed of false alarms; however, although false alarms during periods of especially high dynamic stability were rare, when such false alarms did occur, the pattern of reaction times was consistent with a class of “premeditated” false alarms. We next considered how false alarm behavior would be related to neural activity at the other end of the spectrum, during periods of relatively low dynamic stability.

### Unstable sensory activity near attractor boundaries and “misperception” false alarms

Under the dynamic attractor framework, state instability would be most *critical* when the state is near the attractor boundary. That is, unstable perturbations far from the attractor boundary would be less likely to push the visual system into the false alarm basin of attraction, whereas similar-magnitude perturbations near the boundary are more likely to do so. Such an excursion could be called a “misperception false alarm”, since we are considering activity in a cortical area with a predominantly sensory function. To test this, we sought to compare the strength of the relationship between stability and false alarm rate when the state was (1) near the attractor boundary versus (2) far from the attractor boundary. Because we had already determined that especially high (above-median) stability could be explained as “premeditated” false alarms that were likely due to executive function rather than misinterpretation of sensory signals, we restricted this analysis to trials with below-median stability index, where proximity to attractor boundaries was more likely to be consequential.

Because we do not have explicit knowledge of the dynamic landscape for the system, but can only infer dynamics in the vicinity of states visited by the system by observation, we do not know where the attractor boundaries are. However, we reasoned that the distance of the system state from attractor boundaries likely varies over the course of the trial in a relatively consistent way, which would lead to *some* relationship between SI and time elapsed during the trial (i.e., ISI). Such a relationship could be idiosyncratic for each monkey due to different behavioral strategies or differences in anatomy.

As predicted, we found that the relationship between SI and false alarm rate for both monkeys was not constant over the course of a trial, but rather varied in strength in a consistent way as a function of ISI (Figure 7). This is consistent with the idea that the system approaches and recedes from attractor boundaries over the course of a trial, and dynamic instability is more critical (i.e., makes the difference between committing or avoiding a false alarm) when near such a boundary.

For both monkeys, the relationship between SI and false alarm rate was weaker at shorter ISIs (the 2nd and 3rd *ISI* deciles in Figure 7b; 300 ms was the shortest ISI in the experiment), but then grew to a significantly stronger relationship peaking at intermediate ISIs (5th decile for both monkeys; ∼470 ms for monkey M1 and ∼520 ms for monkey M2). This relationship then weakened again for ISIs in the 6th to 8th decile, before growing to another peak at relatively higher ISIs (9th decile for both monkeys; ∼560 ms for M1, and ∼600 ms for monkey M2). Because the period between peak relationships was approximately 80–90 ms, this could be consistent with the neural state traversing a limit cycle attractor at a frequency around 11–12.5 Hz.

Our earlier analysis found that false alarms following especially stable periods of V4 activity were better explained as “premeditated” false alarms, arising from executive function and not dependent on signals from sensory cortex. In that case, one would predict that the proximity of V4 activity to attractor boundaries should make little difference for premeditated false alarms. To test this, we divided our trials into two groups, one with greater stability than the median (putative “premeditated” trials), and the other group with below median stability (putatively susceptible to “misperception”). As predicted, we found that the modulation of SI-FA relationships with ISI was only observed for the trials putatively susceptible to misperception, and not for the “premeditated” trials (supp. fig. 3).

## Discussion

We asked whether dynamic (in)stability in visual cortical activity could explain animals’ false alarm behavior on a task requiring long periods of vigilant inaction. We found that animals were less likely to commit false alarms following periods of high dynamic stability in V4 and that such dynamic stability also led to a slowing of saccadic actions (both correct and incorrect actions). On a more fine-grained level, our results were consistent with a break-down of false alarms into two broad categories: “premeditated” false alarms, characterized by high dynamic stability in V4 overall, but little dependence on the moment-to-moment details of the activity, and “misperception” false alarms, consistent with perturbations in V4 activity cascading across attractor boundaries during periods of low dynamic stability. Thus we found that activity in the sensory cortex during this task exhibited the characteristics of a dynamical system, and those dynamics were consequential for animal behavior.

Such organization as a complex dynamical system may be an important and universal principle underlying cortical computation. Much of the research into how the dynamical systems framework explains brain circuit function has come through the study of skeletomuscular motor control, but it has been unclear the extent to which the same framework applies across different functional domains. The current findings in the visual cortex would be consistent with a universal principle.

The dynamical systems framework is natural for motor control, where the coordinated kinematics of muscles unfold in time. In this context, motor *preparation* is viewed as setting the initial condition, which allows the appropriate motor action to unfold (Vyas *et al*., 2020). The motor preparatory activity uses mixtures of neurons that are orthogonal to those that drive motor outputs, which enables the preparatory state to be set covertly without leading to premature motor action. We recently showed evidence that a similar principle guides the preparation of visuospatial selective attention in V4 (Snyder *et al*., 2018): a consistent attention-dependent system state is established prior to stimulus onset, but that state does not change the overall level of V4 activity; once the stimulus perturbs the state, however, the response unfolds differentially depending on that covert initial condition. We showed that a minimal dynamical systems model recreated the experimentally observed patterns of neurophysiological results. The dynamics of visual perception may even be directly linked to those of motor control through the process of biological motion perception (Krakowski *et al*., 2011), through a sort of dynamical analog of so-called “mirror neurons” (Rizzolatti & Craighero, 2004) that recognize supramodal features of the dynamics underlying of visual perception of actions and motor execution thereof.

Another computational advantage of dynamical systems is their allowance for pattern completion: partial input that pushes the state into an appropriate basin of attraction is sufficient to lead to the execution of the full pattern of activity. In the motor domain, Li *et al*. (2016) tasked mice to locate objects by whisking, then during movement preparation the experimenters optogenetically suppressed large portions of the premotor cortical network. They found that the premotor network was able to compensate for the missing activity consistent with pattern completion by a dynamic attractor, leading to motor behavior being largely unaffected. For sensory systems, such dynamical pattern completion may be critical to provide stable categorical perception in the face of noisy and dynamic input, such as perceptual “closure” of occluded objects (Doniger *et al*., 2000; Tang *et al*., 2018), and could underlie some illusions, such as illusory contours (Altschuler *et al*., 2012). Aberrations in the dynamic landscapes that support these completion processes may help to explain some disorders of perception, such as sensory hallucinations with psychosis (Waters *et al*., 2014) or dementia (Barnes & David, 2001), and elevated sensitivity in sensory processing disorder or Autism spectrum disorders (Marco *et al*., 2011). On the other end of the spectrum, overly stable dynamics could also be problematic. For example, one interpretation of obsessive-compulsive disorder (OCD) is as a tendency for overly strong attraction of neurophysiological limit cycles guiding behavior (Rolls *et al*., 2008). While high-level behaviors receive much of the attention in the study of OCD, deficits in low-level visual processing and perceptual decision-making have also been reported (Goncalves *et al*., 2010; Kim *et al*., 2008). A general change in dynamic stability could provide a unifying framework for understanding this constellation of symptoms, and point to perceptual assays that could be used as biomarkers for mental disorders.

While much of the study of dynamics in motor and premotor cortex has concerned the planning and execution of movements, there has been recent interest in how premotor dynamics support value-based decision-making. For example, Wang *et al*. (2023) tasked monkeys to pick between offered rewards of different magnitudes and delays, signified by different symbolic visual cues, while the experimenters recorded population activity in lateral prefrontal cortex. Similar to our approach, the researchers estimated the attractive strength of dynamic attractors from the residuals of individual-trial neural population state space trajectories. They found that the strength of dynamic attraction was related to the consistency of animals’ decisions in the task. Our current results in visual cortex are largely consistent with this previous finding in prefrontal cortex, and further enable us to dissect animals’ behavior at a finer-grained scale. Specifically, the stable dynamics underlying consistent decisions that Wang *et al*. (2023) observed could be consistent with what we termed “premeditated” false alarms in this task, as well as confident judgements of correctly detected targets. It is possible that the stable dynamics in visual cortex associated with these premeditated false alarms are inherited from prefrontal feedback. However, we also found evidence that periods of low stability can “make or break” animals perceptual judgements on our task, suggesting a different class of false alarm behavior based in visual misperception.

We found that for trials with relatively low stability (i.e., not likely to result in premeditated false alarms), the relationship between dynamic stability and false alarm rate varied in a reliable way over time since the preceding stimulus. Namely, the relationship started out weak, but grew to a significant peak at regular and repeated intervals (∼10–12.5 Hz; fig. 7b). This periodic relationship could be consistent with the system traversing a limit cycle that approaches an attractor boundary separating perceptual from motoric states. The 10–12.5 Hz frequency we observed corresponds to the so-called “alpha” band that has been linked to attentional suppression of visual processing (Foxe & Snyder, 2011; Snyder & Foxe, 2010; Banerjee *et al*., 2011; Mathewson *et al*., 2011). Neural oscillations have also been implicated in the growing evidence in support of a “rhythmic” theory of attention that holds that theta oscillations organize alternate time periods suitable for perceptual processing and motoric action, and the current results are certainly consistent with this framework (Fiebelkorn *et al*., 2011; Fiebelkorn & Kastner, 2019; Aussel *et al*., 2023).

Taken together, the current results add to the growing appreciation of the computational role for dynamical systems in neuroscience by linking dynamical stability in visual cortex to perceptual decisionmaking behavior. Improved understanding of neurophysiological dynamics will likely be critical for intervening in brain function, such as to treat complex mental illnesses. For example, rather than trying to precisely impose a particular pattern of neural activity on the brain through highly targeted stimulation or inactivation, one feasible approach may be to rather shape the dynamic landscape so that neural activity naturally unfolds along more favorable trajectories. Further, monitoring the dynamic stability of neural activity may enable people to monitor for potential errors of perception or decision-making that are critical for navigating daily life.

## Methods

### Ethical oversight

Experimental procedures were approved by the Institutional Animal Care and Use Committee of the University of Pittsburgh and were performed in accordance with the United States National Research Council’s *Guide for the Care and Use of Laboratory Animals*.

### Subjects

Two adult male rhesus macaques (*Macaca mulatta*) were used for this study. Surgeries were performed in aseptic conditions under isoflurane anesthesia. Opiate analgesics were used to minimize pain and discomfort perioperatively. A titanium head post was attached to the skull with titanium screws to immobilize the head during experiments. After each subject was trained to perform the spatial attention task, we implanted a 96-electrode Utah array (Blackrock Microsystems, Salt Lake City, UT, USA) in V4. The array was implanted in the right hemisphere V4 for Monkey M1, and the left V4 for Monkey M2. A detailed description of these methods and separate analyses of a portion of these data were published previously (Snyder *et al*., 2018; Cowley *et al*., 2020; Snyder *et al*., 2021; Umakantha *et al*., 2021; Johnston *et al*., 2022; Sachse & Snyder, 2023).

### Array recordings

Signals from the arrays were band-pass filtered (0.3 - 7500 Hz), digitized at 1 kHz and amplified by a Grapevine system (Ripple Neuro, Salt Lake City, UT, USA). Signals crossing a threshold (periodically adjusted using a multiple of the root-mean-squared noise) were stored for offline analysis as candidate neural spikes. For this report, we analyze only LFPs; identification of candidate spikes was relevant only for receptive field mapping for stimulus selection. LFPs were low-pass filtered on-line at 250 Hz by the Grapevine amplifier, then resampled off-line to 500 Hz.

### Receptive field (RF) mapping

Before beginning the behavioral task, we mapped the receptive fields (RFs) of the spiking neurons recorded on the V4 arrays by presenting small (≈1^°^) sinusoidal gratings (four orientations) at a grid of positions. We subsequently used Gabor stimuli scaled and positioned to roughly cover the aggregate RF area determined by the responses to the small gratings at the grid of positions. For Monkey M1 this was 7.02^°^ full-width at half-maximum (FWHM) centered 7.02^°^ below and 7.02^°^ to the left of fixation, and for Monkey M2 this was 4.70^°^ FWHM centered 2.35^°^ below and 4.70^°^ to the right of fixation. We next measured tuning curves by presenting gratings at the RF area with four orientations and a variety of spatial and temporal frequencies. For each subject we used full-contrast Gabor stimuli with a temporal and spatial frequency that evoked a robust response from the population overall (i.e., our stimulus was not optimized for any single neuron). For Monkey M1 this was 0.85 cycles/^°^ and 8 cycles/s. For Monkey M2 this was 0.85 cycles/^°^ and 7 cycles/s. For the task, we presented a Gabor stimulus at the estimated RF location, at the mirror-symmetric location in the opposite hemifield, or at both locations simultaneously.

### Behavioral task

Subjects maintained central fixation as sequences of Gabor stimuli were presented in one or both of the visual hemifields, and were rewarded with water or juice for detecting a change in orientation of one of the stimuli in the sequence (the target) and making a saccade to that stimulus (Figure 1a). The probable target location was block-randomized such that 90% of the targets would occur in one hemifield until the subject made 80 correct detections in that block (including cue trials, described below), at which point the probable target location was changed to the opposite hemifield. The fixation point was a 0.6^°^ yellow dot at the center of a flat-screen cathode ray tube monitor positioned 36 cm from the subjects’ eyes. The background of the display was 50% gray. We measured monitor luminance gamma functions by photometer and linearized the relationship between input voltage and output luminance using lookup tables.

We tracked the subjects’ gaze using an infrared eye-tracking system (EyeLink 1000; SR Research, Ottawa, Ontario, Canada). Gaze was monitored online by the experimental control software to ensure fixation within ∼ 1^°^ of the central fixation point throughout each trial. After fixating for a randomly chosen duration of 300 to 500 ms (uniformly distributed), a visual stimulus was presented for 400 ms, or until the subjects’ gaze left the fixation window, whichever came first. If the subject’s eyes left the fixation window and subsequently fixated on a non-target stimulus for at least 50 ms, it was considered a *false alarm*.

For the initial trials within a block, a Gabor stimulus was presented only in the hemifield that was chosen to have a high probability of target occurrence for the block. There were two cue conditions: a cue at the RF location (cue-RF) or a cue in the opposite hemifield (cue-away). These cue trials were to alert the subjects to a change in the probable target location and were excluded from the analysis. The initial cue location was counterbalanced across recording sessions. Once a subject correctly detected five orientation changes during the cue trials, bilateral Gabor stimuli were presented for the remainder of the block.

Each trial consisted of a sequence of 400 ms stimulus presentations separated by 300–500 ms interstimulus intervals (uniformly distributed). Stimulus sequences continued until the subject made an eye movement (data during saccades were excluded from analysis), or a target was presented but the subject did not respond to it within 700 ms (i.e., a *Miss*). For Monkey M1, the average trial duration was 2.44 ± 1.42 s (mean ± SD; N = 33344 trials). For Monkey M2, the average trial duration was 2.44 ± 1.41 s (mean ± SD; N = 31556 trials). For the first presentation in a sequence, the orientation of the stimulus at the cued location was randomly chosen to be 45^°^ or 135^°^, and the orientation of the stimulus in the opposite hemifield, if present, was orthogonal to this. Subsequent stimulus presentations in the sequence each had a fixed probability of containing a target (30% for monkey M1, 40% for monkey M2), i.e., a change in orientation of one of the Gabor stimuli compared to the preceding stimulus presentations in the trial. Because each stimulus (after the first) had a fixed probability of being a target, sequence lengths were roughly exponentially distributed; sequences predominantly had two stimulus presentations (standard then target), and very few sequences had more than four stimulus presentations. To compensate for the reward delay discounting phenomenon and encourage engagement with potentially long trial durations, the number of rewards (fluid drops) given for a correct trial increased with the whole number of seconds (*s*) of trial duration following the formula:

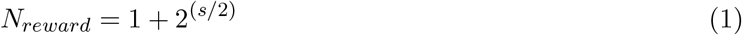

with *N*_*reward*_ ∈ {1, 2, …, 20}. Within a block, 90% of targets (randomly chosen) occurred in one visual hemifield (valid targets) and 10% of targets occurred in the opposite hemifield (invalid targets). For valid targets, the orientation change was randomly chosen to be 1^°^, 3^°^, 6^°^, or 15^°^ in either the clockwise or anti-clockwise direction (monkey M1: 11.49 ± 3.14 (mean ± SD, across sessions) valid targets of each orientation at each location; monkey M2: 14.56 ± 4.75 valid targets of each orientation at each location). For invalid targets, the orientation change was always the near-threshold value of 3^°^, clockwise or anticlockwise (because invalid targets occur infrequently, we restricted the number of orientation change magnitudes for this condition in order to derive a reasonable estimate of the target detection rate). We analyzed trials including either valid or invalid targets, but excluded from analysis all neural data from the time of target onset through the end of the trial. That is, we only analyzed responses to non-targets, of which there were two types: one with a 45^°^ stimulus in the RF, and the other with a 135^°^ stimulus in the RF. Trials where monkeys’ reaction time (square-root transformed) exceeded ±3 standard deviations from the mean, have been excluded from all behavioral analyses (on average 3.27% of all epochs in a session for M1 and 3.19% for M2). For Monkey M1, 1376.58 ± 337.56 (mean ± SD) stimuli with the 45-degree grating in the RF, and 1403.21 ± 325.71 (mean ± SD) stimuli with the 135-degree grating in the RF per session have been included in this study. For Monkey M2, 1812.13 ± 552.87 (mean ± SD) stimuli with the 45-degree grating in the RF, and 1779.13 ± 556.93 (mean ± SD) stimuli with the 135-degree grating in the RF have been included in this study. Monkey M1 completed 25 sessions of the experiment; monkey M2 completed 24 sessions. One session for each subject was subsequently excluded from the analysis because of recording equipment failure.

### Behavioral Analysis

To quantify monkeys’ perceptual sensitivity (*d*^*′*^) in the task, we estimated their *falsealarm* rate (*FA*) and *hit* rate (*Hit*). *Hit* rate was computed as the fraction of *target* presentations, where the monkeys chose the correct *target* location. *FA* rate was computed as the fraction of *non-target* presentations, where the monkeys made incorrect saccades to either of the presented stimuli. We calculated *d*^*′*^ as:

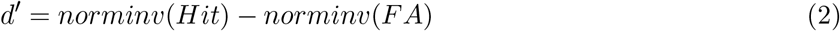

where *norminv* refers to the normal inverse cumulative distribution.

To quantify the association between stimulus repetition number (*repetition* in eq. 3) and *falsealarm* likelihood, we fitted a generalized linear mixed-effect model with the slope of *repetition* (*β* in eq. 4) as the main effect predictor. We used random-effect terms for *intercept* and *slope* of *repetition* grouped by *session* to account for session-specific variations. The Wilkinson notation of the model used is:

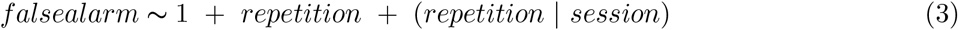

The full regression model can be written as:

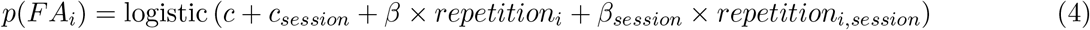

Similarly, we quantified the dependence between stimulus *repetition* and *RT*_*F A*_, using the regression model described in eq. 5a and 5b.

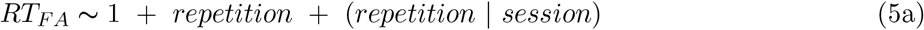

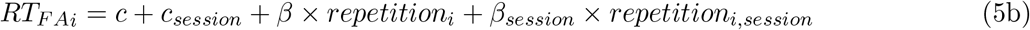

### Data pre-processing

For each session, field potential data were band-pass filtered using a finite-impulse-response (FIR) filter with cut-off frequencies at 1 Hz and 40 Hz. In trials where monkeys correctly withheld saccade, data were epoched for each trial between -300 ms to +700 ms w.r.t. stimulus onset. In trials where monkeys made false alarms, data were epoched both w.r.t. stimulus onset and w.r.t. saccade onset. To detect and exclude artifacts, we computed the peak-to-peak amplitude in the LFPs for each channel on each trial and averaged them across channels to get a single representative value for each trial. Trials where these values exceeded ±4 standard deviations from the mean value were indexed as bad trials and removed from subsequent analysis. For trials with saccades (e.g. *falsealarm* or *correct hits*), we log-transformed the reaction-time values, and trials exceeding ±4 standard deviations from the mean were indexed as outliers and removed from subsequent analyses. After excluding bad trials, we averaged the peak-to-peak amplitudes across trials to get a representative value for each channel. Channels for which these values exceeded ±4 standard deviations from the mean, were indexed as *bad* channels and were interpolated using all other channels weighted by the inverse of the distance between the bad channel and *good* channel, where all the weights summed to a unit.

### Dimensionality Reduction

#### Demixed principal components analysis (dPCA)

To confirm that LFPs reliably encoded relevant task variables such as repetition number, we used dPCA. dPCA decomposes population activity into multiple components that explain as much variance in the data as possible, while maximally capturing the dependence between individual task parameters and neural signals. We used cue, orientation of stimulus in the RF, and stimulus repetition number as the marginalization factors for performing dPCA. Details of this method can be found elsewhere (Kobak *et al*., 2016). We used publicly available MATLAB code to perform this analysis (https://github.com/machenslab/dPCA).

#### Generalized eigenvalue problem (GEV)

GEV, or joint diagonalization, is a general framework underlying various commonly used source separation techniques such as Common Spatial Patterns, Independent Component Analysis, Denoising Source Separation etc. (for reviews see: de Cheveigné & Parra (2014); Särelä *et al*. (2005)). It seeks to extract components from multidimensional data that maximize signal-to-noise ratio. The flexibility and versatility of the tool lie in defining what constitutes the signal in the data. For example, one common problem faced in electrophysiological recordings is the contamination of neural signals by ambient electrical line noise (i.e. 60 Hz for our data). Here, we can use GEV to identify components that maximally explain the 60 Hz signal in the data while minimally capturing signals in other frequency bands which are potentially neural signals. The data then can be denoised by regressing out the contribution of such components.

In our case, we wanted to extract components that maximally capture neural activity common across trials with minimal residual activity. So, we defined signal as the trial-averaged LFPs (*X*_*signal*_ in eq. 6). *X*_*signal*_ has a dimensionality of *T* × *D* (2 ms time bins times the number of channels).

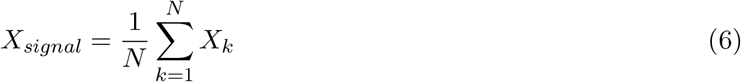

Thereby, we defined *noise* as the residuals after subtracting out the *signal* from the data *X* (eq. 7). *X*_*noise*_ has a dimensionality of *T* × *D* × *N* (time bins times channels times number of trials).

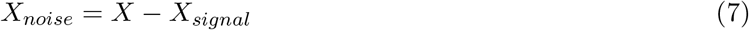

To compute the covariance matrices corresponding to *signal* and *noise*, we estimated *μ*_*signal*_ and *μ*_*noise*_ by averaging *X*_*signal*_ and *X*_*noise*_ along the time-dimension, *T* (eq. 8a and 8b). Both *μ*_*signal*_ and *μ*_*noise*_ are of dimension 1 × *D* each.

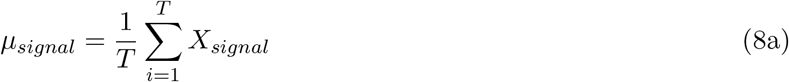

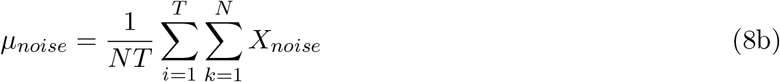

The *signal* covariance and *noise* covariance can be computed as:

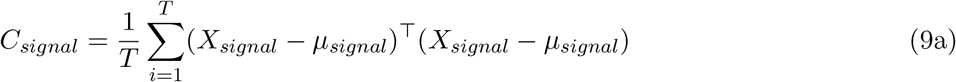

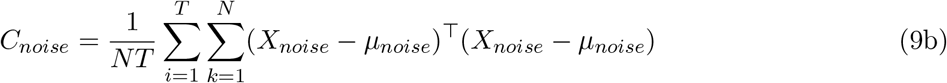

To extract components, i.e. a linear transformation (*W*) when applied to raw data (*X*), that capture maximum signal variance with minimum residual noise variance, we can define the generalized eigenvalue problem as:

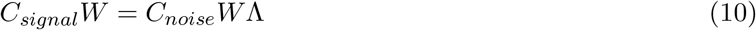

Eq. 10 can be re-written as:

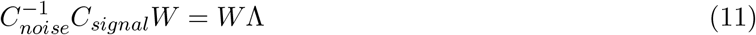

Since in most cases of electrophysiological data, *C*_*noise*_ can be rank-deficient, the computation of *W* can be performed in the following manner:

1. diagonalization of *C*_*noise*_, i.e.

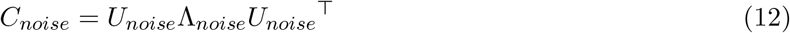
2. After removing the components contributing negligible variance to *C*_*noise*_, *C*_*signal*_ is diagonalized in the following manner:

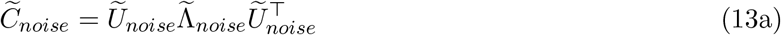

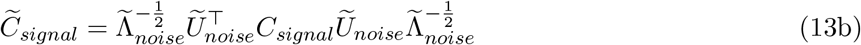
3. *W* is computed as:

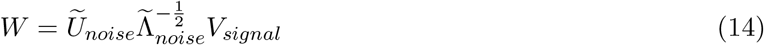

where *V*_*signal*_ are eigenvectors of 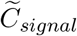 i.e.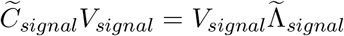.

For each trial *k*, the signal component of *X*_*k*_ that is most consistent (*stable*) across trials, i.e. *S*_*k*_, is estimated by eq. 15a. The *stable* trajectory of *X* is computed as the trial-average of *S*_*k*_ (eq. 15b).

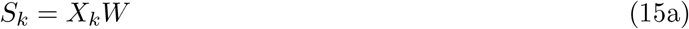

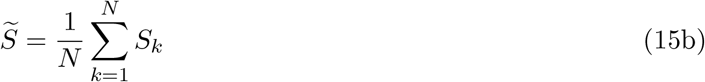

We used the publicly available MATLAB toolbox NoiseTools for this analysis (http://audition.ens.fr/adc/NoiseTools/). The diagonal elements of Λ index the signal-to-noise ratio in each dimension of the multidimensional space. For subsequent analyses, we kept the top dimensions that cumulatively exceed 95% of the matrix trace of Λ (for monkey M1: range 4 − 22 (*median* = 5); and for monkey M2: range 7 − 12 (*median* = 8) dimensions).

### Stability Estimation

One approach in studies of dynamical systems approaches in neuroscience has been to fit parameters of a linear time-invariant dynamical systems model to neural data, and then interpret the fitted parameters. One limitation of this approach is that the strength of conclusions that may be drawn depends on the quality of the fit of the model to the data (i.e., the estimated model parameters are hard to interpret if the model is a poor fit). Moreover, such approaches include assumptions that are likely violated for neural signals (i.e., linearity and time-invariance). Such limitations could have an out-sized influence when aiming to study noisy neurophysiological data at the single-trial level, as we did for this study. To eschew these limitations, we used a data-driven estimate of dynamic stability that does not depend on modeling. We reasoned that a system’s degree of stability is reflected in its response to external perturbations. When the system’s states deviate from its steady trajectory because of perturbation, it should exert force in the direction opposite to the perturbation to bring it back to its steady state. In other words, the *pulling force* opposite to the perturbations signifies the degree of stability.

We conceptualized the residual activity as the perturbations away from the steady trajectory i.e. the trial-averaged neural data, and estimated the moment-by-moment perturbations using eq. 16. For simplicity, only the time dimension has been denoted in the subsequent equations in this section and has been indexed using the subscript *t*.

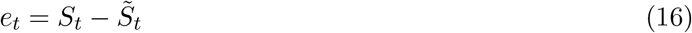

We estimated the moment-by-moment changes in these perturbations using as follows:

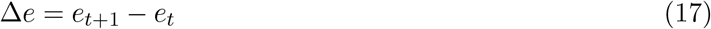

The activity acting against the said perturbations at each time step can be quantified as follows:

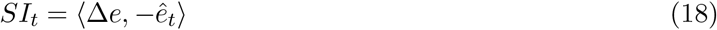

where *SI*_*t*_ refers to estimates of stability index at time *t* and *ê*_*t*_ is the unit norm vector corresponding *e*_*t*_, i.e. 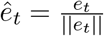.

To normalize the *SI* values, before estimating *SI* we applied a “z-score” transformation to Δ*e* along the *time* dimension in each trial. As the trial-by-trial Δ*e* estimates were very noisy, where appropriate (Figure 3a, and 3c), we applied a 150ms moving-window average to smooth the resultant *SI* estimates. Smaller window sizes produced qualitatively similar results. In analyses concerning epochs where *time* was aligned to stimuli onset (Figure 3a, 3b, and 3c), in each trial we subtracted the average of *SI* within the time window [-200 ms, 0 ms] from the entire epoch as a baseline correction.

### Statistical Analysis

To test the regression coefficients between false alarm rate and repetition number (Figure 1c), we used a two-sided hypothesis test of the t-statistics estimated from the coefficient estimates and corresponding standard errors, with sessions as the degrees of freedom and significance threshold *α* = 0.05. A similar procedure was used to test the regression coefficients between false alarm RT and repetition number (Figure 1d).

Furthermore, we fitted generalized linear mixed-models between false alarm occurrence (1_*F A*_ : 1 if a *falsealarm* occurred, 0 otherwise) and repetition number after combining data from all the sessions for each monkey (section *False alarm behavior improved with repetition number*). We iterated the regression model 1000 times using a bootstrapping procedure where we sampled the observations with replacement in each iteration. We estimated the significance level of the regression coefficients as the percentage of coefficient estimates in the bootstrapped distribution that lied below zero. Similarly, for the generalized linear mixed-model between *RT*_*F A*_ and stimulus *repetition*, We estimated the significance level of the regression coefficients as the percentage of coefficient estimates in the bootstrapped distribution that lied above zero.

To test the decoding performance of LFPs (section *Variance in LFPs explained by repetition number*), we used the *dpca classificationAccuracy* function of the dPCA toolbox. For each session, we organized the LFP data as a 6-D matrix (*Y*) of dimension *C* × *M*_1_ × *M*_2_ × *M*_3_ × *T* × *N*_*M*_, where *C* refers to channel index (96 channels). *M*_1_, *M*_2_ and *M*_3_ refer to number of conditions in the marginalization factors used, i.e. *cue* (*M*_1_ = 2), *orientation* of the stimulus in RF (*M*_2_ = 2) and stimulus *repetition* (*M*_3_ = 4, repetition number ≥ 4 were grouped). *N*_*M*_ is the maximum number of trials corresponding to any condition in any of the marginalization factors. The entries in the matrix where *N*_*M*_ exceeded the number of possible trials for a condition were filled with *NaN* (not-a-number). To estimate the discriminability of different conditions for each marginalization, a sub-matrix *Y* ^*test*^ was carved out of *Y*, consisting of a random combination of trials for each condition and each marginalization. *Y* ^*test*^ was of dimension *C* × *M*_1_ × *M*_2_ × *M*_3_ × *T*. The remainder of *Y* was considered as the training data i.e. *Y* ^*train*^. dPCA was performed on the marginalization average of *Y* ^*train*^, i.e. 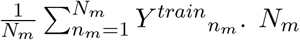. *N*_*m*_ denotes the number of trials for each condition in each marginalization. Both *Y* ^*train*^ and *Y* ^*test*^ were projected onto the dPCA estimated space, producing *Z*^*train*^ and *Z*^*test*^. At each time point *t* and marginalization (*M*_*m*_) in *Y* ^*test*^, classification was done based on the distance of *Z*^*test*^ from the marginalization average of *Z*^*train*^. We iterated this procedure 50 times in each session. We averaged the classification accuracy over the time window [0, 400 ms] w.r.t. stimulus onset to get a representative accuracy value for each session. For each marginalization, we used a *two*-tailed *t*-test to evaluate if the estimated accuracies were significantly different from the corresponding *chance* value, e.g. since there were 4 conditions for stimulus *repetition, chance* accuracy was 25%.

To test the association between *SI* and stimulus repetition number (fig. 3a), we estimated the Spearman’s rank correlation between stimulus repetition number and *SI* at each time point *t* within each session (*ρ* of dimension *T* × *K, K* refers to the number of sessions). Unless otherwise noted, we applied a “Fisher z-transformation” (eq. 19) to correlation values throughout the study, before performing any statistical test on them. We used a cluster-based permutation test to evaluate the statistical significance of *z*_*ρ*_ (Maris & Oostenveld (2007)).

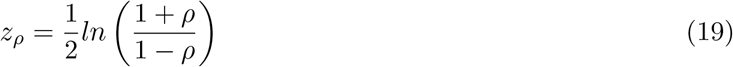

We performed a two-sided t-test at each time point, using *α* = 0.05 as a preliminary significance threshold to test if *z*_*ρ*_ is significantly different from 0. We summed the t-scores of adjacent significant time points (a “cluster”). The resultant sum is the “cluster statistic”. We then randomly permuted the group labels between *z*_*ρ*_ and [0] along the *session* dimension while keeping the *time* dimension consistent between the two. We performed a two-sided t-test between the two pseudo-groups, identified clusters in the permuted groups, and stored only the cluster score with the largest absolute value. We repeated this procedure for 10000 times. The originally identified clusters for which the absolute values of corresponding cluster statistics were above the 95 percentile of the absolute values of the permuted cluster scores were deemed statistically significant.

In Figure 3b, for each trial we computed the average *SI* within the time window [100 ms, 400 ms] w.r.t. stimulus onset. We estimated the Spearman’s rank correlation between this average *SI* and stimuli repetition number within a session. We used a *right* -tailed *sign-rank* test to evaluate if the median correlation was significantly above 0.

In Figure 3c, to further signify the association between repetition number and *SI*, we estimated the rank correlation between repetition number and *SI* across sessions, after averaging the *SI* values for each repetition number within a session (resulting matrix *ρ* is of dimension *T* × 4 × *K*; trials with stimuli *repetition number* ≥ 4 were grouped). To perform a cluster-based permutation test and estimate cluster statistics, we transformed the correlation values to *t* -scores using eq. 20. We permuted the repetition number values across sessions, computed the correlation between the permuted repetition number and *SI*, converted the estimates to t-scores, and estimated the pseudo cluster score. This procedure was repeated 1000 times to get a distribution of pseudo-cluster statistics, against which the statistical significance of the original clusters was tested.

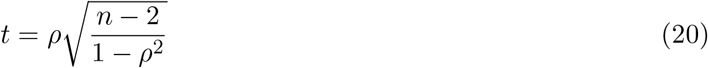

To test the association between *SI* and *RT*_*F A*_ (Figure 4a, and 4b), we computed the partial rank correlation between *SI* and *RT* in each *session* to account for variations due to stimuli repetition number and ISI in *RT*_*F A*_. Similarly, the partial rank correlation was computed between *SI* and *RT*_*hit*_ to account for variations due to stimulus *repetition* and *target* orientation change magnitude. In Figure 4a and 4c, cluster-based permutation tests were used to identify clusters along the *time* dimension, for which the estimated correlations were significantly different from 0 across *sessions* (1000 permutations). In Figure 4b and 4d, we computed the partial correlation between *RT* and *SI* estimates averaged within the time window [-600 ms, -100 ms] w.r.t. saccade onset in each trial. We performed a *two*-tailed *t*-test to test statistical significance.

**Figure 4.**
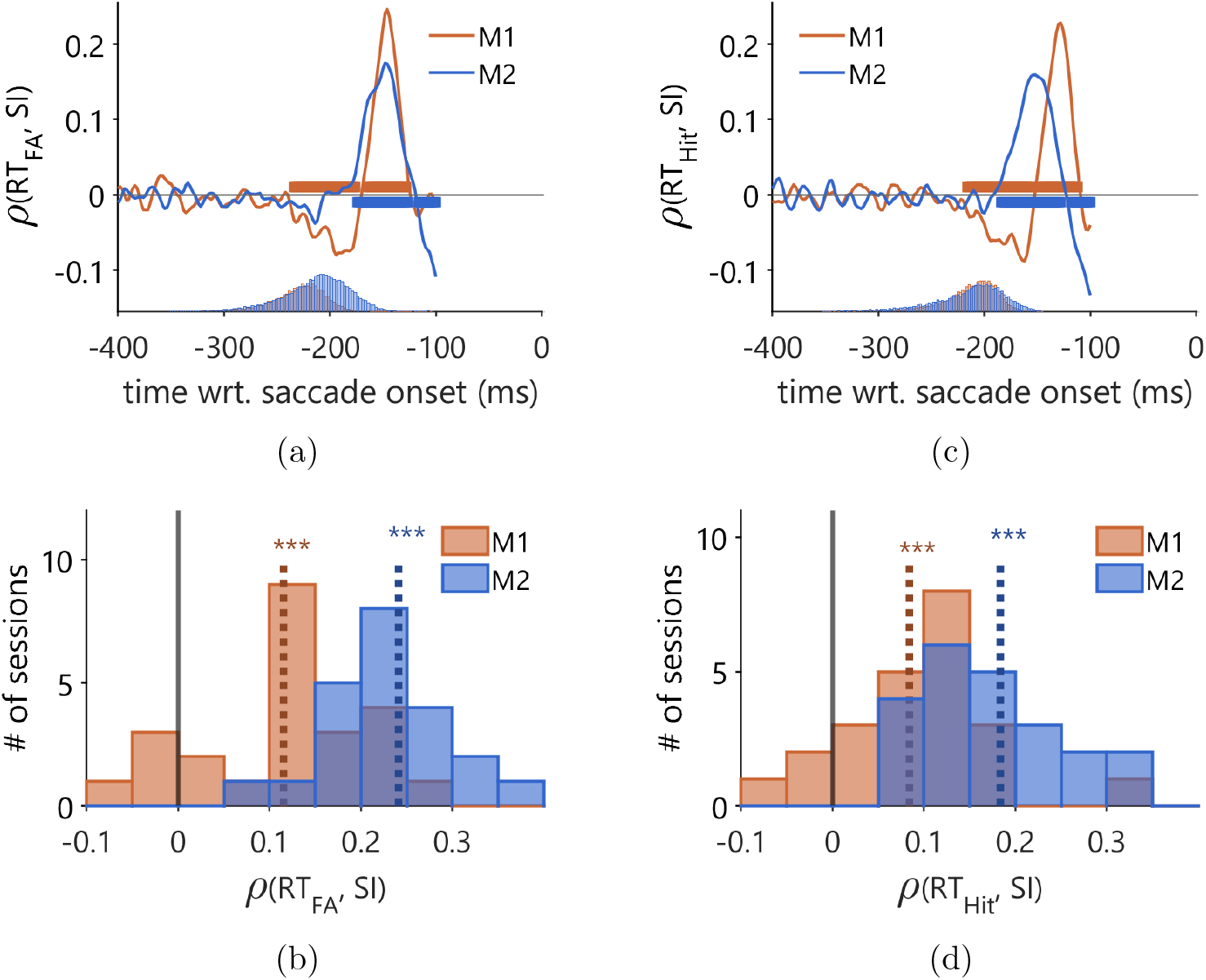
Reaction time for both false alarm and correct saccades was slower when neural activity preceding them was more dynamically stable. (a) Time series of partial correlation between stability and *RT*_*F A*_, i.e. *ρ*(*RT*_*F A*_, *SI*) accounting for stimulus repetition number, estimated using signal aligned to the saccade onset. The inline histograms show relative stimulus onset w.r.t. the saccade onset for the two monkeys. Because stimulus onset relative to saccade time was variable, data on each trial from the time of stimulus onset through the saccade was excluded from this analysis. (b) Histogram of partial *ρ*(*RT*_*F A*_, *SI*) accounting for stimulus repetition number and inter-stimulus intervals (ISIs), across sessions for the two monkeys, showing an increase in *RT*_*F A*_ with *SI*. The stability index was averaged over a time window [-600 ms, -100 ms] relative to saccade onset. The dotted lines indicate the average correlation value across sessions. Statistics were performed using two-tailed t-test (*** : *p <* 0.001). (c) Time series plot of partial correlation between *SI* and *RT* for the correct hits, after accounting for stimuli repetition number and target change amplitude. (d) Similar to (b), but for correct hits. Both (b) and (d) show that in general, *RT* increases with an increase in the stability of the neural trajectories.

To test the association between GFP and *SI* (Figure 5b), for each session, we averaged the GFP within a *SI* quintile and normalized the *session* average to unit-value (resultant GFP matrix is of dimension *T* × 5 × *K*). At each time point *t*, we computed the rank correlation between GFP and *SI* bin numbers across sessions and used a cluster-based permutation test as described above in this section, to identify statistically significant clusters along the *time* dimension, where the estimated correlations were significantly different from 0 (1000 permutations).

**Figure 5.**
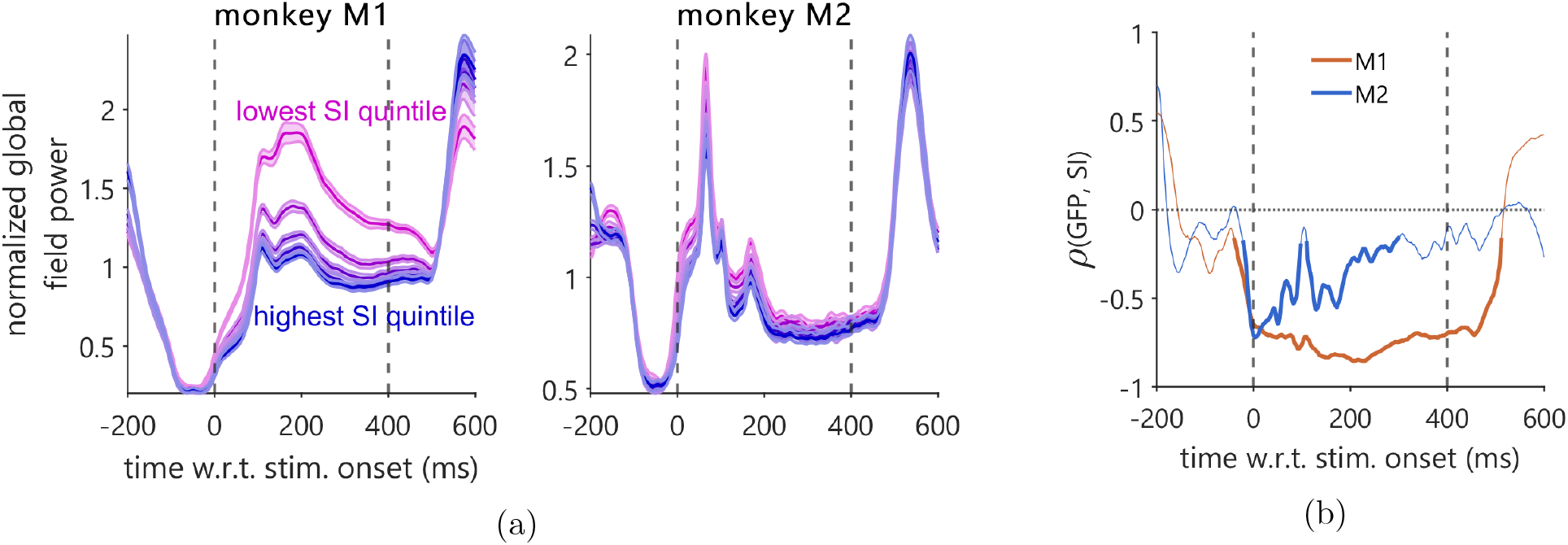
Relationship between stability and global field power (GFP). (a) Time series of normalized (relative to session-average) GFP averaged across sessions for individual monkeys. Each line corresponds to GFPs averaged across trials in a quintile of *SI* estimates. Blue and magenta lines correspond to the highest and the lowest quintiles of SI estimates, respectively. (b) Rank correlation between stability index (SI) bin numbers and average GFP for each bin, estimated across sessions for individual monkeys. Thick lines denote statistically significant correlation values (two-tailed cluster-based permutation test, *p* ≤ 0.05).

To test the differences in *RT*_*F A*_ (Figure 6b) between different quadrants outlined in Figure 6a, we used a *two*-tailed *t*-test with the number of observations in each quadrant corresponding to degrees of freedom.

**Figure 6.**
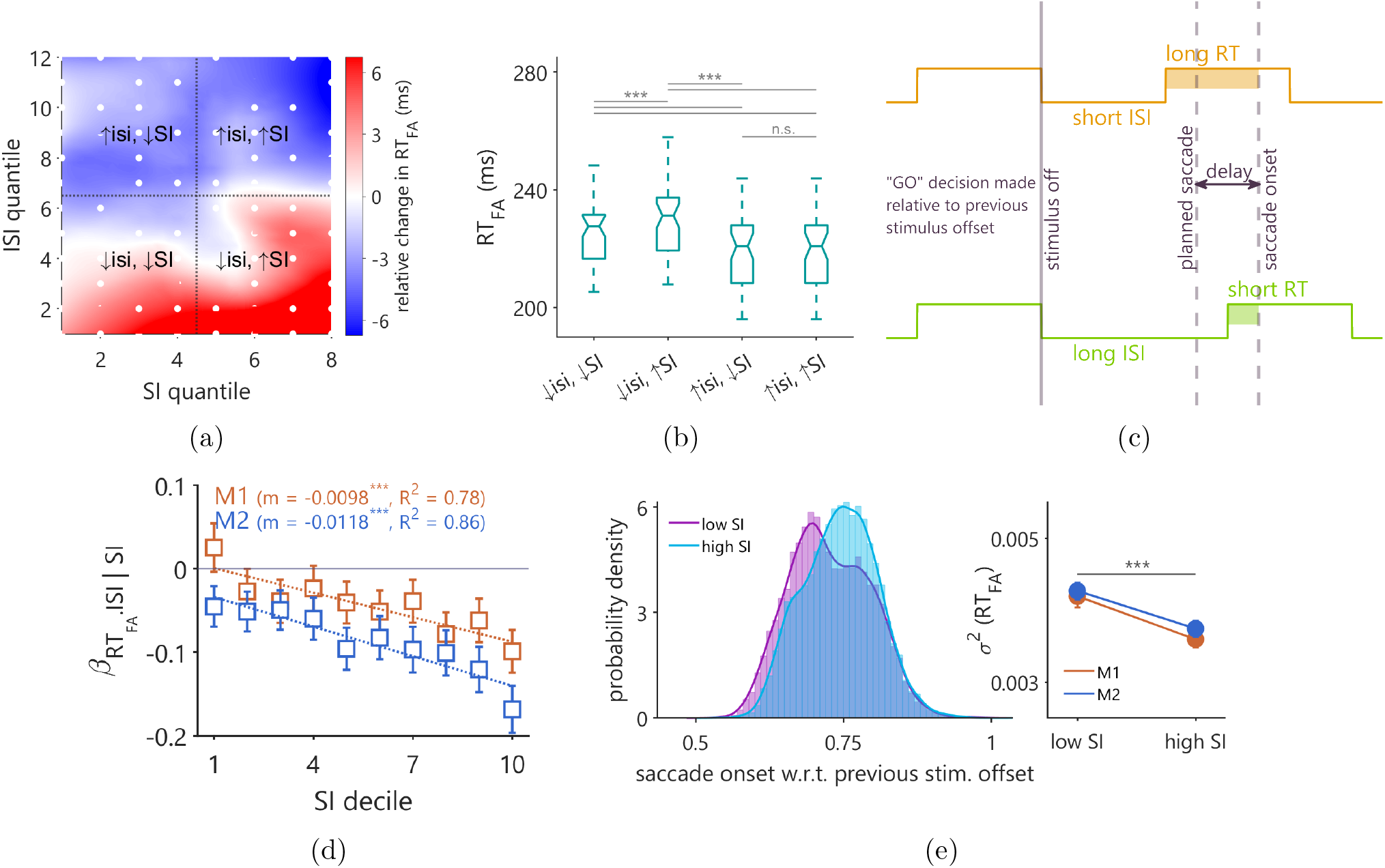
Variation in *RT*_*F A*_ with *ISI* during planned false alarm saccades. (a) 2D distribution of relative changes in *RT*_*F A*_ for different (*SI, ISI*) bins, showing a relatively greater rate of change in *RT*_*F A*_ with *ISI* for higher *SI* compared to that for lower *SI*. (a) Similar 2D distribution for change in %*FA* for different (*SI, ISI*) bins. It shows an increase in FA rate with *ISI* only for *SI* bins of lower magnitude. (b) Boxplot of *RT*_*F A*_ for the four combinations of SI and ISI i.e., (low ISI, low SI), (lowISI, high SI), (high ISI, low SI) and (high ISI, high SI). (b) Same conventions as (b) for *FA* rate. (c) Schematic describing a negative relationship between RT and ISI for planned saccades. (d) Plot of the slope between *RT*_*F A*_ and *ISI* for each *SI* decile, after partialing out the trend between *ISI* vs. *SI*, and, *RT*_*F A*_ vs. *SI*. Errorbars correspond to 95% CI calculated as 1.96 × *SE*. The plot shows the slope becoming steeper with an increase in *SI*. (e) Left: histogram of time duration between saccade onset and previous stimulus offset for low and high *SI*; right: plot of variance of the distributions shown on the left. Errorbars correspond to 95% CI of the variance estimates from the bootstrapped distributions.

**Figure 7.**
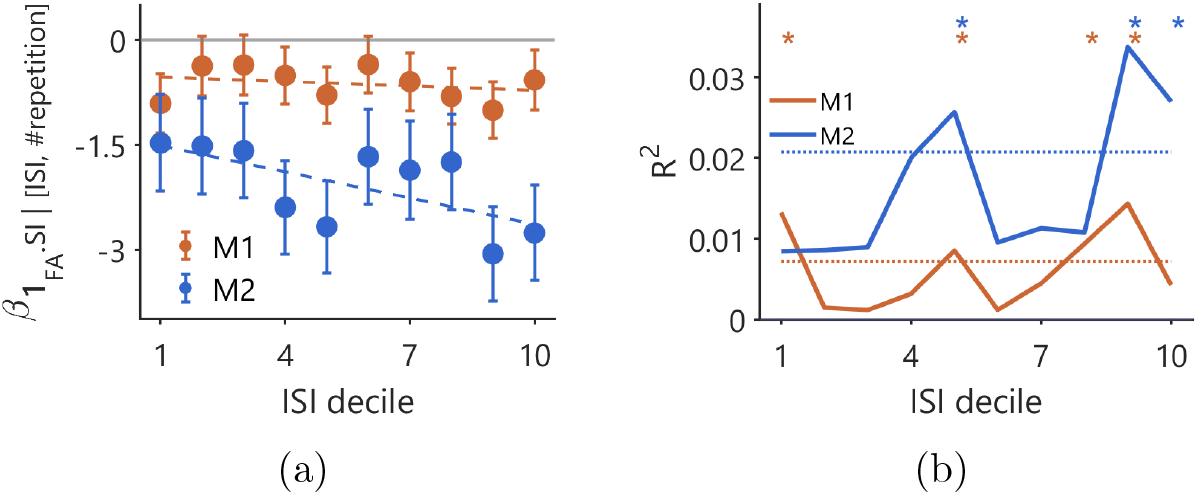
Variation in *FA* likelihood with *SI* for different *ISI* bins. (a) Plot of coefficients of regression between false alarm occurrence (𝟙_*F A*_: indicator function to denote false alarm occurrence) and stability for different *ISI* bins, after regressing out variation in individual factors contributed by *ISI* and stimuli repetition number. (b) Fraction of variance in false alarm occurrence explained by SI for different ISI bins. Thick lines denote estimated *R*^2^ from the regression between false alarm occurrence and SI and dotted lines denote 95 percentile estimates of *R*^2^ after permuting trial labels corresponding ISI bins. *: ISI bins for which original *R*^2^ values were significantly larger (*p <* 0.05, one-tailed) compared to the permuted ones.

To test the trend in the slope 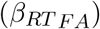 between *RT*_*F A*_ and ISI with *SI* decile index ([1,2,…,10]), we fitted a linear regression model between the two (Figure 6d). *p*-value was estimated from the F-statistics corresponding to the regression coefficients. The *R*^2^ value denoted the goodness-of-fit of the regression.

To test the differences between the variances (*σ*^2^(*RT*_*F A*_)) in saccade times aligned to the preceding stimulus offset, corresponding to *low* and *high SI* values, we used a bootstrapping procedure to create a distribution of estimates of *σ*^2^(*RT*_*F A*_). We sampled the 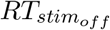 distribution with replacement and estimated its variance. This was performed 1000 times. We used a two-tailed t-test to determine statistical significance between the bootstrapped distributions of *σ*^2^ corresponding to *low* and *high* SI.

To observe the trend in the explaining power of *SI* for false alarm occurrence (𝟙_*F A*_), as a function of ISI, we used the following procedure. For trials corresponding to each ISI decile, we fitted a GLM between 𝟙_*F A*_ and *SI*, after regressing out contributions of ISIs within the decile considered and stimuli repetition number to 𝟙_*F A*_. We computed the *R*^2^ adjusted for the number of coefficients. We sampled 10% of the trials at random without replacement and estimated the *R*^2^ for this subset of trials. We performed this operation 20000 times to create a null distribution of *R*^2^, that ignored any systemic trend w.r.t. ISI. ISI indices where the original *R*^2^ exceeded 95 percentile estimates of this null distribution were marked statistically significant.

## Acknowledgements

A.C.S. was supported by NIH grants K99/R00EY025768, R01EY028811 and R01EY011749; a NARSAD Young Investigator award from the Brain & Behavior Research Foundation; and an Alfred P. Sloan Foundation research fellowship. Experimentation was performed by A.C.S. while a postdoctoral fellow in the laboratory of Matthew A. Smith at the University of Pittsburgh (now at Carnegie Mellon University). We thank Dr. Smith for his support, including funding (NIH grants R01MH118929, R01EB026953, R01EY022928 and P30EY008098; NSF NCS BCS 1954107/1734916; Research to Prevent Blindness; and the Eye and Ear Foundation of Pittsburgh). The authors would like to thank Ms. Samantha Schmitt for assistance with surgery and data collection, and Dinah McAlly for feedback on dynamical systems analyses.

## Author Contributions

B.S.: Methodology, Software, Formal Analysis, Writing - Original Draft, Writing - Review and Editing, Visualization. A.C.S.: Conceptualization, Methodology, Software, Formal Analysis, Investigation, Resources, Data Curation, Writing - Original Draft, Writing - Review and Editing, Visualization, Supervision, Project Administration, Funding Acquisition.

---

**Supplemental Figure 1:**
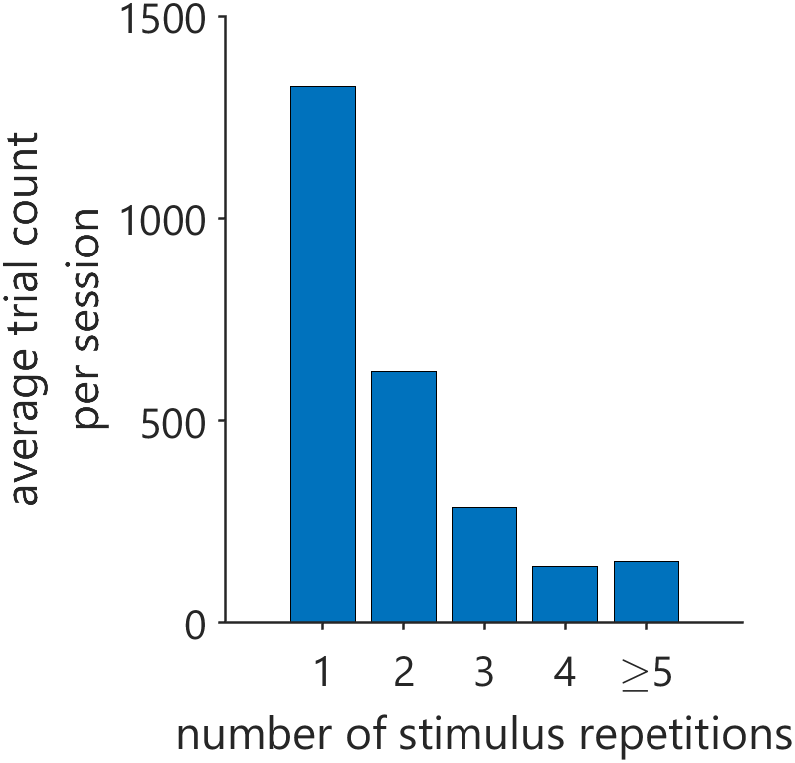
Histogram showing trial counts in a session where stimuli presentation was repeated *N* number of times, where *N* ∈ {1, 2, 3, 4, ≥ 5}

**Supplemental Figure 2:**
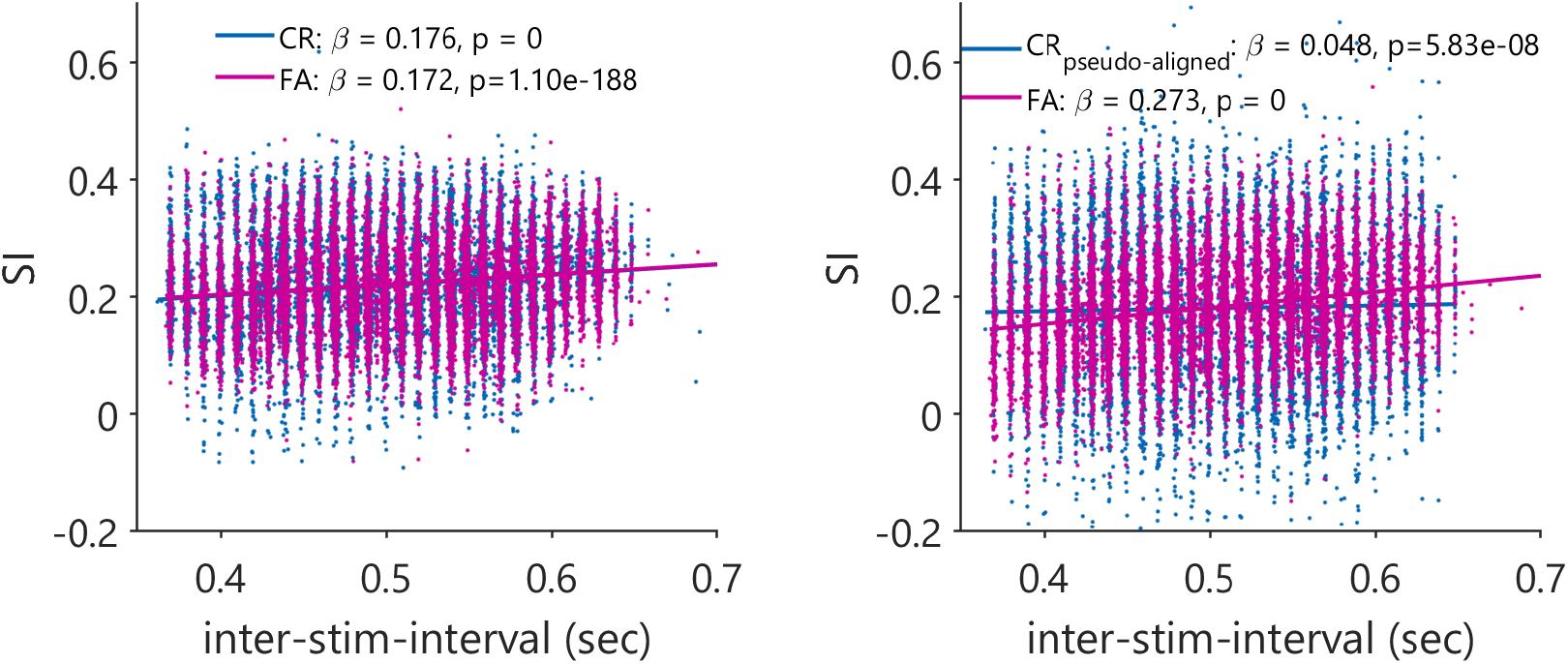
Scatter plot showing a positive relationship between *SI* and *ISI*. CR: correct-reject; FA: false alarm. CR_pseudo-aligned_: correct-rejection trial subset selected to match ISI distribution of the FA trials.

**Supplemental Figure 3:**
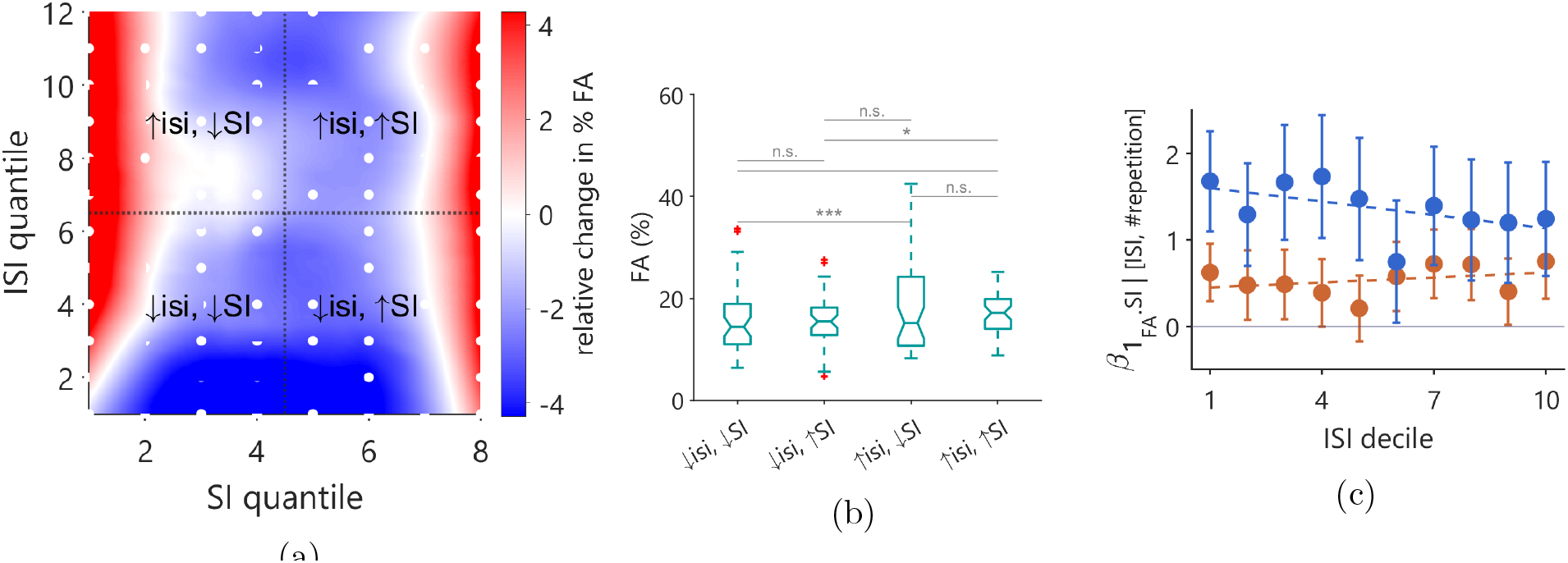
2D Distribution of relative changes in *FA*% with SI and ISI. (a) Relative change in *FA*% for different (*SI, ISI*) bins. (b) Average *FA*% for the four quadrants partitioned according low and high ISIs and SI. (c) Regression coefficient between FA occurrence and SI, for high *SI*.

